# Deep metabolic profiling assessment of tissue extraction protocols for three model organisms

**DOI:** 10.1101/2021.12.16.472947

**Authors:** Hagen M. Gegner, Nils Mechtel, Elena Heidenreich, Angela Wirth, Fabiola Garcia Cortizo, Katrin Bennewitz, Thomas Fleming, Carolin Andresen, Marc Freichel, Aurelio Teleman, Jens Kroll, Rüdiger Hell, Gernot Poschet

## Abstract

Metabolic profiling harbors the potential to better understand various disease entities such as cancer, diabetes, Alzheimer’s, Parkinson’s disease or COVID-19. Deciphering these intricate pathways in human studies requires large sample sizes as a means of reducing variability. While such broad human studies have discovered new associations between a given disease and certain affected metabolites, i.e. biomarkers, they often provide limited functional insights. To design more standardized experiments, reduce variability in the measurements and better resolve the functional component of such dynamic metabolic profiles, model organisms are frequently used. Standardized rearing conditions and uniform sampling strategies are prerequisites towards a successful metabolomic study. However, further aspects such as the choice of extraction protocol and analytical technique can influence the outcome drastically. Here, we employed a highly standardized metabolic profiling assay analyzing 630 metabolites across three commonly used model organisms (Drosophila, mouse and Zebrafish) to find the optimal extraction protocols for various matrices. Focusing on parameters such as metabolite coverage, metabolite yield and variance between replicates we compared seven extraction protocols. We found that the application of a combination of 75% ethanol and methyl tertiary-butyl ether (MTBE), while not producing the broadest coverage and highest yields, was the most reproducible extraction protocol. We were able to determine up to 530 metabolites in mouse kidney samples, 509 in mouse liver, 422 in Zebrafish and 388 in Drosophila and discovered a core overlap of 261 metabolites in these four matrices. To enable other scientists to search for the most suitable extraction protocol in their experimental context and interact with this comprehensive data, we have integrated our data set in the open-source shiny app “MetaboExtract”. This will enable scientists to search for their metabolite or metabolite class of interest, compare it across the different tested extraction protocols and sample types as well as find reference concentrations.

## Introduction

Metabolomics, defined as the separation and subsequent measurement of small molecules in either qualitative or quantitative way, enables the generation of metabolic profiles of any sample of interest. While genomics and transcriptomics are analyzed within the framework of a single organism and understood by the blueprint of genes or transcripts of the respective species, metabolomics encompasses all compounds that may be metabolized by an organism, its microbiome or that are introduced by the environment (‘exposome’) at a given time. Therefore, the metabolome incorporates the environmental influence as well as interactions with other organisms (Johnson et al., 2012). It can serve as a bridge between the organism, its interactions and any disease, e.g. between diet, the gut microbiome and metabolic disease (Pallister et al., 2017). While the complexity and dynamic nature of the metabolome is daunting from an analytical perspective, metabolomics harbors the potential to better understand as well as diagnose various disease entities such as diabetes (Arneth et al., 2019), kidney disease (Abbiss et al., 2019), Parkinson’s (Shao & Le, 2019), Alzheimer’s disease (Wilkins & Trushina, 2018) and most recently, COVID-19 (Sindelar et al., 2021).

The potential to understand the metabolic signatures of any given disease entity is tremendous, however, deciphering the intricate underpinnings of those in a human study requires costly, as well as time and work extensive population-wide association studies with several hundred participants per group (Nicholson et al., 2008). These broad studies may be successful in the discovery of new associations between a respective disease and the measured metabolites, i.e., biomarkers, but they are limited in their mechanistic explanations despite all efforts. While the dynamic nature of the metabolome provides incredibly powerful insights, it also highlights the challenges of metabolome analyses - its variability and associated pitfalls.

Variation and noise that are biologically inherent or are introduced at some point to the sample are complicating metabolic analyses, impairing the quality of the findings, limiting their validity and may even overshadow the effect size of the research question itself. Factors that introduce such variability range from intrinsic ones derived from the study organism, e.g. age or sex (Bell et al., 2021; Brennan & Gibbons, 2020), to extrinsic factors, e.g. diet, lifestyle or medication (Adamski, 2016; Mellert et al., 2011). Additionally, other factors, such as pre-analytical ones during sample collection (Lippi et al., 2020; Yin et al., 2015), or analytical factors deriving from the sample preparation, the extraction protocols or analytical approach used to conduct the measurement (Erben et al., 2021; Lin et al., 2007) are also influential and need to be accounted for.

Model organisms that are reared under controlled laboratory conditions and manipulated genetically to analyze a certain genotype address several of the challenges mentioned above. Combining the standardized rearing conditions and stringent sampling protocols with the already extensive knowledge accumulated from other “-omics” in models such as mice, Drosophila or Zebrafish enhances the explanatory power of metabolomic studies tremendously while simultaneously reducing the number of samples needed to generate meaningful results. To ensure that the analytical phase, i.e. the extraction and measurement of metabolites, does not introduce biases and variability, an in-depth evaluation of such aspects is necessary.

Standardized metabolomic analyses are commercially available by companies such as Metabolon (www.metabolon.com) or Biocrates (www.biocrates.com). The latter has developed standardized and robust LC-MS/MS based kits which enable the absolute quantification of specific compound classes or more broadly, up to 630 metabolites in the case of the MxP Quant 500 kit (Biocrates). Within these 630 metabolites, the MxP Quant 500 kit covers 14 small molecule and 9 different lipid classes. Due to its standardized nature and compatibility with a multitude of LC-MS/MS platforms, data generated via such a kit-based approach enables inter-and intra-laboratory comparability (Siskos et al., 2017), as well as its integration from different experiments. Although these kits were initially developed for human biofluids, i.e. plasma and serum, they may be used for tissue samples and other sample types such as cultured cells (Andresen et al., 2021) or supernatants likewise. However, there is no consensus on the optimal metabolite extraction procedure amongst the metabolomic community for the investigation of polar and non-polar metabolites covering that many chemical classes within one analysis.

In this study, this open question was addressed using the highly standardized targeted metabolomics kit (Biocrates MxP Quant 500) to evaluate seven extraction protocols designed to extract both, polar and non-polar metabolites, differing in their solvent composition and extraction mode (mono/bi-phasic) as well as handling complexity (Figure 1.). We compared the metabolite coverage, concentration yields and robustness, i.e. the coefficient of variance (CV%) across three commonly used model organisms (mice, Zebrafish and Drosophila), focusing on either whole organisms as sample type (Drosophila) or specific organs (liver and kidney) of the respective model organism. Lastly, we integrated our data in the Shiny app “MetaboExtract” (Andresen et al., 2021) to provide a useful source of metabolite concentrations across model organisms and enable other scientists to search for an optimal extraction procedure for their metabolite or metabolite class of interest.

**Figure 1.**
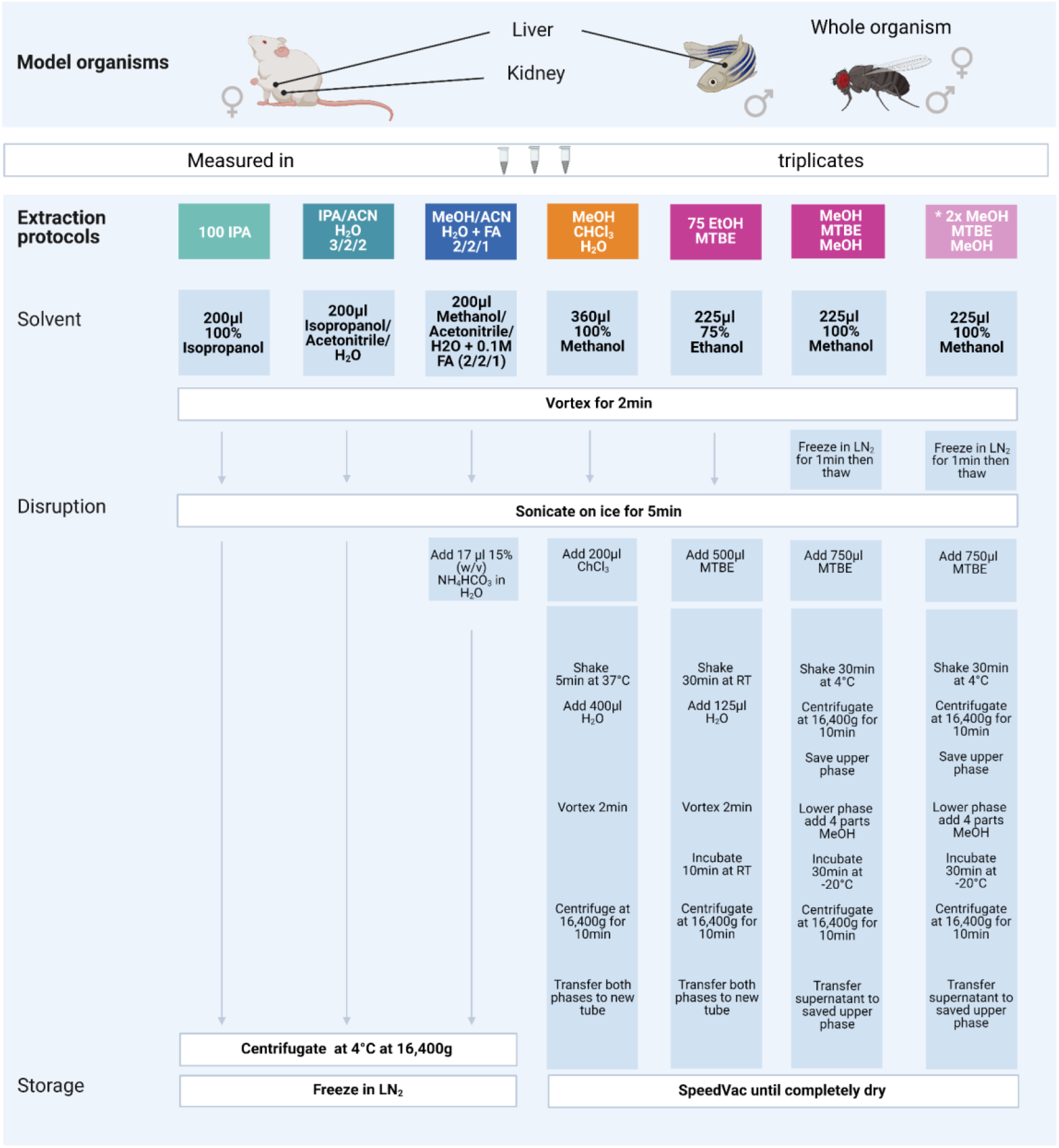
Overview of the seven extraction protocols used as well as the model organisms and sample types investigated. The protocols increase in handling effort and complexity from left to right. The color code indicates similarities amongst the protocols either through solvents or chemicals used. All extraction products were stored at -80°C until further processing. A list of abbreviations can be found above.

## Materials and Methods

### Chemicals

Chemicals were bought from Sigma-Aldrich (Germany). All solvents used for sample extractions and LC-or FIA-MS/MS analyses were of UHPLC-MS quality.

### Model organism growth/culturing conditions

#### Mouse – *Mus musculus*

Nine-week-old C57Bl6N wildtype mice (Charles River, Germany) were anesthetized with isoflurane and blood was taken to generate EDTA-Plasma. Without regaining consciousness mice were killed by cervical dislocation. Livers and kidneys were excised rapidly and shortly rinsed in ice-cold 0.9 % NaCl. Excess liquid was removed before whole organ weight was determined for later normalization and tissue was snap-frozen in liquid nitrogen. All procedures were approved by the Animal Care and Use Committee at the Regierungspräsidium Karlsruhe, Germany (T-40/20).

#### Fly - *Drosophila melanogaster*

W1118 flies were acquired from Bloomington Drosophila Stock Center. For all metabolic measurements, flies were grown under controlled conditions: Flies were allowed to lay eggs on apple plates for 12 hours. First instar larvae hatching within a 4- or 6-hours window were picked and seeded at a density of 60 animals per vial. Adult flies of all genotypes enclosing within a 24-hour time-window were separated by gender in groups of 30 flies and aged for 10 days. Flies were grown and maintained on food consisting of the following ingredients for 30 liters of food: 480g agar, 660g sugar syrup, 2400g malt, 2400g corn meal, 300g soymeal, 540g yeast, 72g nipagin, 187mL propionic acid and 18.7 mL phosphoric acid. At sample collection, flies were pooled and snap-frozen for metabolic profiling.

#### Zebrafish - *Danio rerio*

Adult Zebrafish were kept under a 13-hour light / 11-hour dark cycle and feeding of Zebrafish took places twice a day, freshly hatched *Artemia salina* in the morning and fish flake food in the afternoon. All experimental interventions on animals were approved by the local government authority, Regierungspräsidium Karlsruhe and by Medical Faculty Mannheim (I-19/02) and carried out in accordance with the approved guidelines. Age of adult male Zebrafish was 9 months and both sexes were included. Zebrafish were sacrificed in ice water and livers were immediately dissected and frozen in liquid nitrogen and subsequently stored at -80°C. 7 mg to 10 mg of livers were used for further analysis.

### Sample preparation

To ensure sufficient input material across the model organisms we used 30 pooled flies (w1118 flies), 7-10 mg of Zebrafish liver or 20-22 mg of mouse (C57Bl6N) liver and kidney pooled from three individuals respectively. All tissue samples were pulverized using a ball mill (MM400, Retsch) with precooled beakers and stainless-steel balls for 30 seconds at the highest frequency (30 Hz). The exact weight was determined for normalization of all measurements.

### Metabolite extraction protocols

Here we evaluated six different extraction protocols that are described in Figure 1. We developed these protocols based on own preliminary experience and scanning of general metabolomics literature. The protocol “*MeOH/MTBE*”, noted with an asterisk, was applied twice for mouse samples only. We are including this variation as an additional method (*2xMeOH/MTBE*) when we are referring to the seven extraction protocols.

Briefly, pulverized and frozen samples were extracted using the indicated solvents and subsequent steps of the respective protocol (Fig.1.). After a final centrifugation step the solvent extract of the protocols *100IPA, IPA/ACN* and *MeOH/ACN* were transferred into a new 1.5ml tube (Eppendorf) and snap-frozen until kit preparation. The remaining protocols were dried using an Eppendorf Concentrator Plus set to no heat, stored at -80°C and reconstituted in 120 µl isopropanol (60 µl of 100% isopropanol, followed by 60 µl of 30% isopropanol in water) before the measurement.

### Standardized targeted metabolic profiling

After conducting the described seven extraction protocols, tissue extracts were processed following the manufacturer’s protocol of the MxP® Quant 500 kit (Biocrates). Briefly, 10 µl of the samples or blanks were pipetted on the 96 well-plate based kit containing calibrators and internal standards using an automated liquid handling station (epMotion 5075, Eppendorf) and subsequently dried under a nitrogen stream using a positive pressure manifold (Waters). Afterwards, 50 µl phenyl isothiocyanate 5% (PITC) was added to each well to derivatize amino acids and biogenic amines. After 1 h incubation time at room temperature, the plate was dried again. To resolve all extracted metabolites 300 µl of 5 mM ammonium acetate in methanol were pipetted to each filter and incubated for 30 min. The extract was eluted into a new 96-well plate using positive pressure. For the LC-MS/MS analyses 150 µl of the extract was diluted with an equal volume of water. Similarly, for the FIA-MS/MS analyses 10 µl extract was diluted with 490 µl of FIA solvent (provided by Biocrates). After dilution, LC-MS/MS and FIA-MS/MS measurements were performed in positive and negative mode. For chromatographic separation an UPLC I-class PLUS (Waters) system was used coupled to a SCIEX QTRAP 6500+ mass spectrometry system in electrospray ionization (ESI) mode. LC gradient composition and specific 50×2.1mm column are provided by Biocrates. Data was recorded using the Analyst (Version 1.7.2 Sciex) software suite and further processed via Met*IDQ* software (Oxygen-DB110-3005). All metabolites were identified using isotopically labeled internal standards and multiple reaction monitoring (MRM) using optimized MS conditions as provided by Biocrates. For quantification either a seven-point calibration curve or one-point calibration was used depending on the metabolite class.

## Data processing and analyses

### Validation and filtering

Data validation and quantification was performed using MetIDQ (Oxygen-DB110-3005). Here, metabolites were further categorized based on their quantitation ranges. Additional filtering per metabolite was based on the limit of detection (LOD), limit of quantification (LOQ) and concentration within the quantitative range (valid). To remove metabolites that were not present in any model organism and sample type, we considered a metabolite as detectable when at least 2 out of 3 replicates within a tested protocol were above LOD (see Figure 2.). These metabolites are also visualized as Venn diagrams in Figure 5 for the extraction protocol EtOH/MTBE and for the remaining extraction protocols in supplementary Figure S5. An overview of the LOD, LOQ and valid metabolite proportions are shown in supplementary Figure S1. For all detectable metabolites, the coefficient of variation (CV) in percentage was calculated as well as the median absolute deviation (MAD) based on the concentrations.

**Figure 2.**
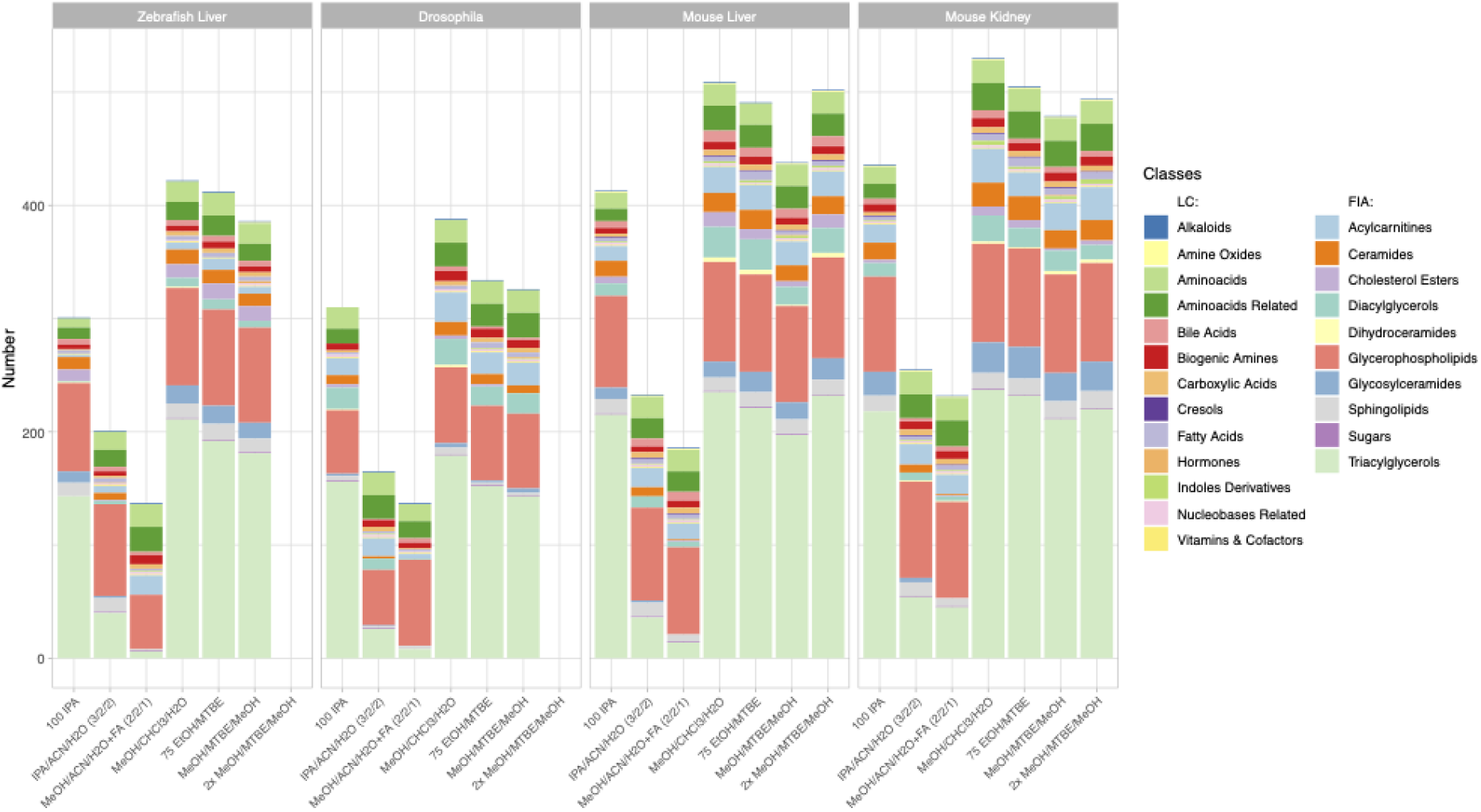
Metabolite coverage per extraction protocol across all sample types and model organisms. Indicated by color are the different metabolite classes measured. A metabolite was counted as detectable when at least 2 out of 3 replicates were >LOD within a given extraction protocol. The legend is categorized between compound classes measured via LC-MS/MS or FIA-MS/MS. Ordering from left to right follows the level of complexity and required time per extraction.

### Statistical analysis

To find the optimal protocol per model organism and sample type, we analyzed the yield per metabolite achieved across the extraction protocols. For this comparison, missing values and zero values were imputed per metabolite with 20% of the minimal positive value of a given metabolite. Subsequently, to perform statistical analyses, the data was log2-transformed. We then performed an ANOVA per metabolite considering all metabolites that were detectable with at least a single method. Extraction protocols were used as categorical variables and concentration as dependent variables. A Tukey post-hoc test (*alpha* = 0.05) was used to determine the extraction protocols with the highest median yield as well as non-significantly (p-adjusted > 0.05) lower yields. These extraction protocols were considered optimal, counted and depicted in Figure 3. Conversely, metabolites that were significantly better extracted in a single extraction protocol are depicted in supplementary Figure S4.

**Figure 3.**
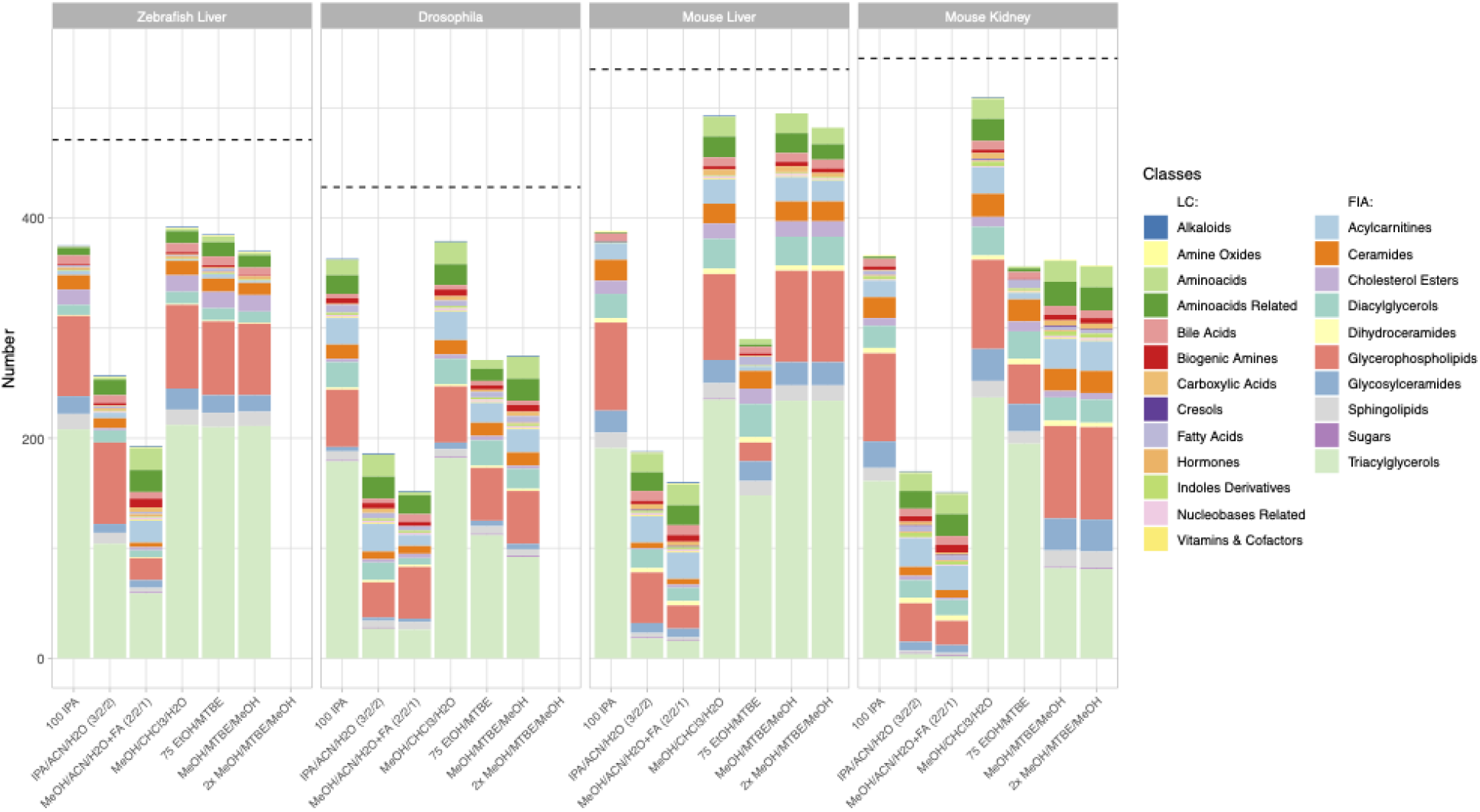
Number of metabolites per class with the highest yield per extraction protocol across all sample types and model organisms. Metabolites that appear in the bar chart are only counted when they produce the highest or a non-significantly lower concentration than another tested extraction protocol (see material and method part). The dotted line shows the number of detectable metabolites for each sample type. Indicated by color are the different metabolite classes measured. The legend is categorized between the LC-MS/MS and FIA-MS/MS measurements. Of note, the metabolite classes that were best suited for a single extraction protocol are depicted in supplementary Figure S4.

### R Shiny app

Data can be explored and downloaded using the Shiny app “MetaboExtract” which is available at http://www.metaboextract.shiny.dkfz.de. The underlying code is also available at https://github.com/andresenc/MetaboExtract (Andresen et al., 2021). Figure 2 and 3 as well as S3 were extracted from the Shiny app.

## Results

The aim of this study was the comparison of seven extraction protocols (Figure 1) across three model organisms to determine the optimal extraction procedure with regards to metabolite coverage, yield and robustness (CV%). In total, we analyzed 630 metabolites, however, after filtering for low concentrated metabolites below the limit of detection (LOD), we continued the analyses using this processed data. Figure S1. shows the ratio of LOD, LOQ and valid measurements per extraction protocol across all sample types and model organisms.

### Biphasic extractions generate the highest coverage and concentration

The metabolic profiling kit (Biocrates MxP Quant 500) used for this study quantifies polar as well as non-polar metabolites across 14 small molecule and 9 different lipid classes. Therefore, an extraction procedure is required that enables solubilization ranging from very polar metabolites (e.g. carbohydrates and amino acids) to very non-polar metabolites such as triacylglycerols (Figure 2.). While the maximum coverage between the different model organisms is expected to be variable, the general performance of the respective protocol remained similar. Figure 2. shows the detected metabolites per extraction protocol across all sample types and model organisms. The ordering of the protocols from left to right also indicates the level of complexity and time required for the protocol (Figure 1).

Clear performance trends between the monophasic (*100IPA, IPA/ACN/H2O, MeOH/ACN/H2+FA*) and biphasic (*MeOH/CHl3/H2O, 75EtOH/MTBE, (2x)MeOH/MTBE*) extractions were apparent. The protocol using *MeOH/CHl3/H2O* resulted in the highest metabolite coverage in all sample types and across all organisms (Zebrafish liver (422), Drosophila (388), mouse liver (509), mouse kidney (530)). Similarly, *75EtOH/MTBE*, as well as both *MeOH/MTBE* protocols, produced a broad coverage across all metabolite classes. In other words, all bi-phasic extractions performed well and were comparable regarding their metabolite coverage.

While *IPA*, a rapid and easy to perform single solvent extraction protocol, produced fair coverage, the remaining protocols, both containing acetonitrile, achieved the lowest coverage regardless of the sample type or organism. Comparison of the different metabolite classes reveal that these monophasic extraction protocols failed to extract several lipids, such as di-and triacylglycerols as well as ceramides or cholesterol esters.

Although the coverage of a given extraction is essential, the yield, as the calculated concentration per metabolite, needs to be maximized and loss during the protocol should be avoided. To scrutinize the extraction protocols regarding this criterion we performed an ANOVA (see material and method part), counting the metabolites that reached the highest or a non-significantly lower concentration in a given extraction protocol per model organism (Fig. 3). Therefore, lower counts in Figure 3. indicate that other protocols did extract significantly higher concentrations.

Here, a similar pattern compared to the metabolite coverage (Figure 2) emerged. The protocol using *MeOH/CHl3/H2O* resulted in the highest concentrations of metabolites measured within each metabolite class across all organisms (Zebrafish liver (392), Drosophila (379), mouse liver (493), mouse kidney (510)). Again, all biphasic extraction protocols performed well, apart from *75EtOH/MTBE* in mouse liver, showing significantly lower yields per metabolite than the other protocols with a strong reduction in glycophospholipids. Surprisingly, the rapid *100IPA* protocol produced comparable or higher concentrations than the two MTBE protocols. Within the group of MTBE protocols the combination with EtOH was superior to a single MeOH extraction. Only the *2xMeOH/MTBE* extraction that was applied twice resulted in higher concentrations. Similarly to the coverage of metabolites, both acetonitrile-containing protocols performed worse across all model organisms and sample types.

### Extraction protocol variability as an essential quality parameter

While coverage and yield are important to determine the optimal extraction protocol for the broadest range of metabolite classes, the variability or the coefficient of variance (CV%) of each metabolite between the analysis of biological triplicates informs about the robustness of a protocol. To better understand the variability across the different protocols and compare it to the coverage simultaneously we plotted both as a spider plot in Figure. 4. There, the variability of the measurements between the triplicates per metabolite in CV% ranges from 0-10% (=excellent), 11-20% (=good), 21-30% (=acceptable) and >30% (=not acceptable). The percentage ranges were calculated from the total of 630 possible metabolites, elucidating the variability of a method but also the number of detectable metabolites per method. For example, in mouse kidney, *75EtOH/MTBE* results in 202 (32.1%) metabolites with an excellent CV, 252 (40%) metabolites with a good CV, 21 (3.3%) metabolites with a CV that is acceptable CV and finally, 30 (4.8%) metabolites with a high CV that is not acceptable. Similarly, *100IPA* appears as a well performing choice in this sample type with most measurements in a CV% range from 0-10% (=excellent).

**Figure 4.**
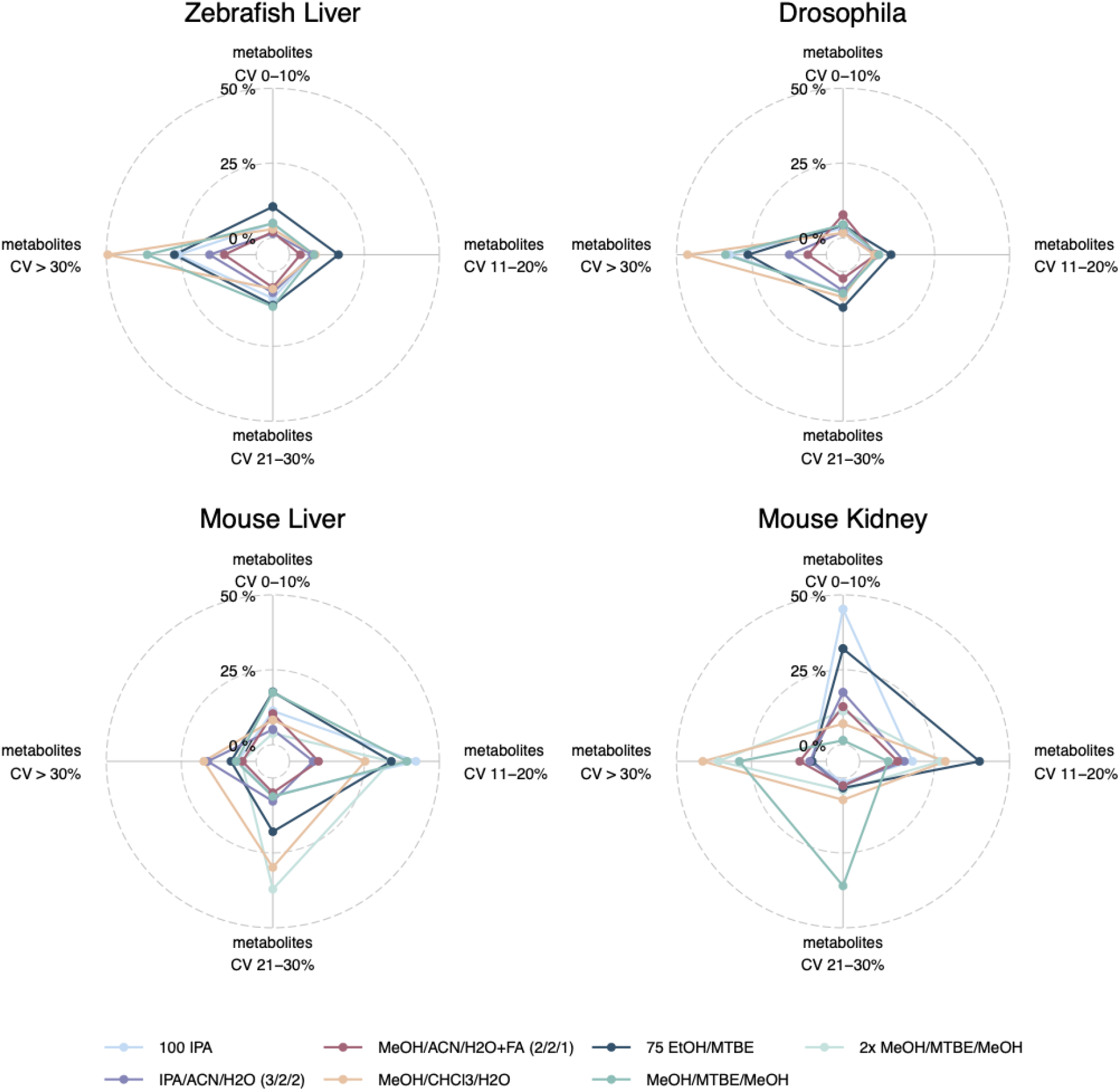
Variability of extraction procedures across all sample types and model organisms. Indicated by color are the different extraction protocols used. The CV% was generated based on the triplicate analyses. For each of the CV% categories the percentage of the total 630 metabolites. Note that *2xMeOH/MTBE/MeOH* was only used for mice sample types. Alternative visualizations of the CV% and MAD are shown in the supplementary figures (Figure S3 and S4).

**Figure 5.**
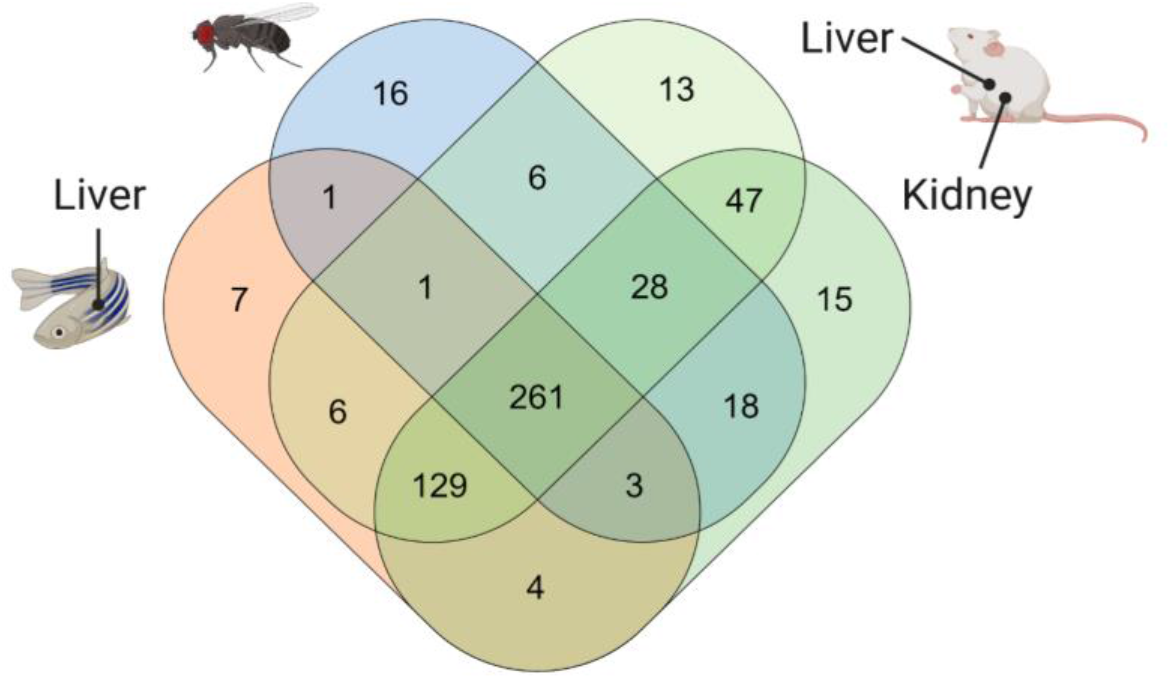
Venn diagram showing number of metabolites above LOD that are common or unique across all sample types and model organisms within the *75EtOH/MTBE* extraction protocol. A comparison of the remaining extraction protocols can be found in the supplement (see supplementary figure S5). A list of the metabolites that are unique and overlapping can be found in the supplementary data file (see supplementary data file S1).

This visualization enables the comparison of several extraction protocols across all sample types regarding their robustness and coverage at once (Figure 4.). The protocol using *MeOH/CHl3/H2O* which resulted in the highest coverage and yield performed the worst across all sample types and model organisms with most of the metabolites with a CV of >30% (Zebrafish liver (312), Drosophila (292), mouse liver (110), mouse kidney (260)). Similarly, *2xMeOH/MTBE* generates a high variability but also a high coverage as well as yield in mouse sample types. Amongst the other model organisms (Drosophila and Zebrafish), a single extraction with *MeOH/MTBE* resulted in a high portion of CV >30% (=not acceptable) compared to *75EtOH/MTBE*. Yet, differences could be seen in mouse sample types, where *75EtOH/MTBE* performed better in mouse kidney than mouse liver and conversely, for *MeOH/MTBE*.

*75EtOH/MTBE* resulted in acceptable levels of variance across all sample types and model organisms (<30%). An alternative visualization of the CV% across the different extraction protocols can be found in supplementary Figure S2. The median and median absolute deviation (MAD) of the coefficient of variation (CV) across the seven extraction protocols is depicted in the supplementary Figure S3. Both visualizations strengthen the conclusion described above.

### Strong overlap of metabolites between the sample types using *75EtOH/MTBE*

Overall, the *75EtOH/MTBE* protocol resulted in the best coverage and highest metabolite concentrations of the evaluated protocols while also showing excellent to acceptable levels of variance between the measurements. As a next step, we used this extraction protocol as an example to visualize the overlap and the uniquely extracted metabolites across the different sample types and model organisms. 261 common metabolites out of 630 possible metabolites can be extracted across all matrices. Additional 126 metabolites are shared between the liver and kidney samples. Only very few metabolites are unique to each sample type, highlighting the broad coverage of the *75EtOH/MTBE* extraction protocol. Venn diagrams for the remaining extraction protocols are provided in supplementary figure S5 as well as the list of shared or unique metabolites in supplementary data file S1. There, *MeOH/CHl3/H2O* is once more the protocol producing the highest coverage across all sample types and model organism (304 metabolites).

### “MetaboExtract” is an interactive resource to explore metabolite extractions and baseline concentrations of model organisms

The presented data provides an attempt to inform about the optimal extraction protocol for a standardized profiling assay. However, it harbors further information such as baseline concentration for sample types and whole model organisms. To access this information researchers may explore the data set via the easy-to-use interactive R/shiny app “MetaboExtract” (Andresen et al., 2021). There, the already present metabolite data on human tissue and cells was expanded by our data set focused on model organisms. Since all data was generated using the standardized MxP Quant 500 assay it is highly comparable. Organisms, tissues, extraction methods and classes of metabolites may be (de)selected to focus on the data of interest that are then provided in comprehensive and interactive visualizations. The data presented in Figure 2 and in supplementary Figure S4 were generated using MetaboExtract. Its standardized nature provides the potential for further expansion via additional MxP Quant 500 assay measurements.

## Discussion

Model organisms enable standardized laboratory-controlled handling, sampling and experiments. This level of standardization in the pre-analytical phase benefits metabolic profiling due to the dynamic nature of the metabolome, rapid turnover rates of metabolites and the influence of the environment (Edison et al., 2016; Saoi & Britz-mckibbin, 2021).

Here, we focused on the extraction and processing of diverse sample types, one of the most important aspects in the analytical phase requiring strict standardization for reproducibility of data. To this end, we used a targeted metabolic profiling approach quantifying up to 630 metabolites across three commonly used model organisms (Drosophila, mouse and Zebrafish) to find a robust, easy-to-use extraction protocol yielding a comprehensive coverage of the target metabolites. This targeted metabolic profiling assay (Biocrates MxP Quant 500) potentially quantifies polar as well as non-polar metabolites across 14 hydrophilic and 9 different lipid classes. Hence, it requires the extraction of a chemically diverse range of metabolites. Besides broad metabolite coverage and high yields in concentrations with little loss during the extraction, we evaluated the robustness of a given method as well as the practicability and effort of performing the protocols.

### Biphasic extractions are superior to monophasic extractions

In our study, biphasic extractions (*MeOH/CHl3/H2O, 75EtOH/MTBE, (2x)MeOH/MTBE*) resulted in superior coverage and concentrations across all model organisms and sample types. Here, the complementary phases, composed of an organic lipid-rich phase and an aqueous phase containing primary and secondary metabolites, were combined and dried in the final step of each protocol allowing for a greater coverage as compared to mono-phasic extractions. Chloroform based bi-phasic extractions by Bligh & Dyer (1959) have been dominantly used over the years due to the focus on the lipid fraction, however, MTBE (methyl tert-butyl ether, i.e. TBME) is more frequently used as a non-toxic and non-carcinogenic alternative to chloroform (Furse et al., 2015; Matyash et al., 2008). Here, both strong hydrophobic solvents in combination with another organic solvent of lower hydrophobicity such as ethanol or methanol resulted in comparable metabolite coverage. Importantly, the chloroform-based extraction resulted in the highest yield as well as broadest coverage, however, substituting it with MTBE resulted in similar but less variable measurements (Figure 3 and 4).

Mono-phasic extractions (*100IPA, IPA/ACN* and *MeOH/ACN*) require less solvents and are performed more rapidly as compared to bi-phasic ones, which is a big advantage when processing large numbers of samples. While the robustness of monophasic extractions was comparable to that of the other well performing biphasic extractions, i.e., *75EtOH/MTBE*, they provided lower compound coverage due to the lack of certain lipids that were poorly extracted, with the exception of *100IPA*, which provided adequate coverage and concentrations in most cases. This easy-to-use and rapid protocol achieved adequate lipid coverage and reproducibility in most model organisms and sample types. However, concentration and coverage for amino acids and amino acid related metabolites were lower as compared to biphasic extraction. Nevertheless, several other lipidomic studies concluded that isopropanol is an adequate alternative to more complex and time-consuming biphasic extractions. There, utilizing isopropanol in a ratio to water, e.g. 90:10 v/v or 75:25 v/v, performed well and was regarded as excellent alternative for lipidomic analyses (Calderón et al., 2019). Although such single phase extractions are in general well suited for lipidomic approaches, the broad nature of the standardized metabolic profiling assay requires a trade-off between coverage, yield as well as reproducibility across all metabolite classes. The latter criterion was recently highlighted by Ghorasaini et al. (2021) in an interlaboratory assessment of extraction protocols for lipidomic analyses. The authors showed that the extraction with MeOH/MTBE performed better and was more practical than the comparable Bligh & Dyer extraction.

In line with this notion and matching the discussed criteria, we recommend the protocol *75EtOH/MTBE* as a suitable broadly applicable biphasic extraction. Importantly, similar conclusions have been made in other sample types. Erben et al. (2021) compared several extraction protocols for metabolic profiling of human stool samples via MxP Quant 500 and Andresen et al. (2021) of human cells from four different tissues (liver and bone marrow) or cell lines (adherent: HEK and non-adherent: HL60), concluded that protocols including methanol or ethanol with MTBE are suitable for these sample types as well, confirming our findings.

### No one size fits all approach

The biphasic MTBE extraction with 75% ethanol (*75EtOH/MTBE*) achieves a broad coverage, high concentrations and little variability in between extractions. These attributes make it a versatile extraction method suitable for the different model organisms and sample types tested.

However, the extraction protocol of choice depends highly on the target as well as the sample type. While our study aimed to find the most versatile protocol it also showed that other protocols extracted certain metabolite classes more efficiently than the broader biphasic extractions. For instance, *MeOH/ACN/H2O+FA* was superior in the extraction of amino acids in Zebrafish and of glycerophospholipids in Drosophila as compared to the other protocols (supplementary Figure S4). Hence, the most suitable extraction protocol highly depends on the type of analysis, i.e., quantitative or qualitative assessment, the coverage needed, i.e., a small set of targets within one metabolite class or a broad screening, as well as the sample type.

To find the most adequate protocol for a given scenario we included this data in a publicly available shiny app - “MetaboExtract” (Andresen et al., 2021). This open access resource is expandable and makes use of the comparability of standardized assays such as MxP Quant 500. MetaboExtract enables users to review and explore standardized extractions and infer baseline concentrations of metabolites across a variety of sample types and organisms. There, users can search for a metabolite or metabolite class of interest, review or compare measured concentrations following a variety of mono- or biphasic extraction protocols across human cells, cell lines and tissue and, now, model organisms.

## Code availability

The custom R scripts (R version 4.0.4) that were used for analysis and visualization are accessible at https://github.com/nilsmechtel. The underlying code for “MetaboExtract” is available at https://github.com/andresenc/MetaboExtract (Andresen et al., 2021).

## List of Abbreviations

LC-MS/MS: Liquid chromatography tandem mass spectrometry
FIA-MS/MS: Flow injection analysis and tandem mass spectrometry
ESI: Electrospray ionization
MRM: Multiple reaction monitoring
EtOH: Ethanol
MeOH: Methanol
MTBE: Methyl tert-butyl ether
ACN: Acetonitrile
IPA: Isopropanol
FA: Formic acid
PITC: Phenylisothiocyanate (or Edman’s Reagent)
LN2: Liquid nitrogen
LOD: Limit of detection
LOQ: Limit of quantification
CV%: Coefficient of variation in percentage
ANOVA: Analysis of variance
MAD: Median absolute deviation

## Acknowledgements

The authors gratefully acknowledge the data storage service SDS@hd supported by the Ministry of Science, Research and the Arts Baden-Württemberg (MWK) and the German Research Foundation (DFG) through grant INST 35/1314-1 FUGG and INST 35/1503-1 FUGG. Figure 1 was created using BioRender.com.

## References

Abbiss, H., Maker, G. L., & Trengove, R. D. (2019). Metabolomics approaches for the diagnosis and understanding of kidney diseases. Metabolites, 9(2). https://doi.org/10.3390/metabo9020034

Adamski, J. (2016). Key elements of metabolomics in the study of biomarkers of diabetes. In Diabetologia (Vol. 59, Issue 12, pp. 2497–2502). Springer Verlag. https://doi.org/10.1007/s00125-016-4044-y

Andresen, C., Boch, T., Gegner, H. M., Mechtel, N., Narr, A., Birgin, E., Rasbach, E., Rahbari, N., Trumpp, A., Poschet, G., Hübschmann, D. (2021). Comparison of extraction methods for intracellular metabolomics. https://doi.org/10.1101/2021.12.15.470649

Arneth, B., Arneth, R., & Shams, M. (2019). Metabolomics of type 1 and type 2 diabetes. International Journal of Molecular Sciences, 20(10), 1–14. https://doi.org/10.3390/ijms20102467

Bell, J. A., Santos Ferreira, D. L., Fraser, A., Soares, A. L. G., Howe, L. D., Lawlor, D. A., Carslake, D., Davey Smith, G., & O’Keeffe, L. M. (2021). Sex differences in systemic metabolites at four life stages: cohort study with repeated metabolomics. BMC Medicine, 19(1), 1–13. https://doi.org/10.1186/s12916-021-01929-2

Bligh, E.G. and Dyer, W. J. (1959). Canadian Journal of Biochemistry and Physiology. Canadian Journal of Biochemistry and Physiology, 37(8).

Brennan, L., & Gibbons, H. (2020). Sex matters: a focus on the impact of biological sex on metabolomic profiles and dietary interventions. Proceedings of the Nutrition Society, 79(2), 205–209. https://doi.org/10.1017/S002966511900106X

Calderón, C., Sanwald, C., Schlotterbeck, J., Drotleff, B., & Lämmerhofer, M. (2019). Comparison of simple monophasic versus classical biphasic extraction protocols for comprehensive UHPLC-MS/MS lipidomic analysis of Hela cells. Analytica Chimica Acta, 1048, 66–74. https://doi.org/10.1016/j.aca.2018.10.035

Edison, A. S., Hall, R. D., Junot, C., Karp, P. D., Kurland, I. J., Mistrik, R., Reed, L. K., Saito, K., Salek, R. M., Steinbeck, C., Sumner, L. W., & Viant, M. R. (2016). The time is right to focus on model organism metabolomes. Metabolites, 6(1). https://doi.org/10.3390/metabo6010008

Erben, V., Poschet, G., Schrotz-King, P., & Brenner, H. (2021). Evaluation of different stool extraction methods for metabolomics measurements in human faecal samples. BMJ Nutrition, Prevention & Health, bmjnph-2020-000202. https://doi.org/10.1136/bmjnph-2020-000202

Furse, S., Egmond, M. R., & Killian, J. A. (2015). Isolation of lipids from biological samples. Molecular Membrane Biology, 32(3), 55–64. https://doi.org/10.3109/09687688.2015.1050468

Ghorasaini, M., Mohammed, Y., Adamski, J., Bettcher, L., Bowden, J. A., Cabruja, M., Contrepois, K., Ellenberger, M., Gajera, B., Haid, M., Hornburg, D., Hunter, C., Jones, C. M., Klein, T., Mayboroda, O., Mirzaian, M., Moaddel, R., Ferrucci, L., Lovett, J., … Giera, M. (2021). Cross-Laboratory Standardization of Preclinical Lipidomics Using Differential Mobility Spectrometry and Multiple Reaction Monitoring. Analytical Chemistry. https://doi.org/10.1021/acs.analchem.1c02826

Johnson, C. H., Patterson, A. D., Idle, J. R., & Gonzalez, F. J. (2012). Xenobiotic metabolomics: Major impact on the metabolome. Annual Review of Pharmacology and Toxicology, 52, 37–56. https://doi.org/10.1146/annurev-pharmtox-010611-134748

Lin, C. Y., Wu, H., Tjeerdema, R. S., & Viant, M. R. (2007). Evaluation of metabolite extraction strategies from tissue samples using NMR metabolomics. Metabolomics, 3(1), 55–67. https://doi.org/10.1007/s11306-006-0043-1

Lippi, G., Von Meyer, A., Cadamuro, J., & Simundic, A. M. (2020). PREDICT: A checklist for preventing preanalytical diagnostic errors in clinical trials. Clinical Chemistry and Laboratory Medicine, 58(4), 518–526. https://doi.org/10.1515/cclm-2019-1089

Matyash, V., Liebisch, G., Kurzchalia, T. V., Shevchenko, A., & Schwudke, D. (2008). Lipid extraction by methyl-terf-butyl ether for high-throughput lipidomics. Journal of Lipid Research, 49(5), 1137–1146. https://doi.org/10.1194/jlr.D700041-JLR200

Mellert, W., Kapp, M., Strauss, V., Wiemer, J., Kamp, H., Walk, T., Looser, R., Prokoudine, A., Fabian, E., Krennrich, G., Herold, M., & van Ravenzwaay, B. (2011). Nutritional impact on the plasma metabolome of rats. Toxicology Letters, 207(2), 173–181. https://doi.org/10.1016/j.toxlet.2011.08.013

Nicholson, J. K., Holmes, E., & Elliott, P. (2008). The metabolome-wide association study: A new look at human disease risk factors. Journal of Proteome Research, 7(9), 3637–3638. https://doi.org/10.1021/pr8005099

Pallister, T., Jackson, M. A., Martin, T. C., Glastonbury, C. A., Jennings, A., Beaumont, M., Mohney, R. P., Small, K. S., MacGregor, A., Steves, C. J., Cassidy, A., Spector, T. D., Menni, C., & Valdes, A. M. (2017). Untangling the relationship between diet and visceral fat mass through blood metabolomics and gut microbiome profiling. International Journal of Obesity, 41(7), 1106–1113. https://doi.org/10.1038/ijo.2017.70

Saoi, M., & Britz-mckibbin, P. (2021). New Advances in Tissue Metabolomics : A Review.

Shao, Y., & Le, W. (2019). Recent advances and perspectives of metabolomics-based investigations in Parkinson’s disease. Molecular Neurodegeneration, 14(1), 1–12. https://doi.org/10.1186/s13024-018-0304-2

Sindelar, M., Stancliffe, E., Schwaiger-Haber, M., Anbukumar, D. S., Adkins-Travis, K., Goss, C. W., O’Halloran, J. A., Mudd, P. A., Liu, W. C., Albrecht, R. A., García-Sastre, A., Shriver, L. P., & Patti, G. J. (2021). Longitudinal metabolomics of human plasma reveals prognostic markers of COVID-19 disease severity. Cell Reports Medicine, 2(8). https://doi.org/10.1016/j.xcrm.2021.100369

Siskos, A. P., Jain, P., Römisch-Margl, W., Bennett, M., Achaintre, D., Asad, Y., Marney, L., Richardson, L., Koulman, A., Griffin, J. L., Raynaud, F., Scalbert, A., Adamski, J., Prehn, C., & Keun, H. C. (2017). Interlaboratory Reproducibility of a Targeted Metabolomics Platform for Analysis of Human Serum and Plasma. Analytical Chemistry, 89(1), 656–665. https://doi.org/10.1021/acs.analchem.6b02930

Wilkins, J. M., & Trushina, E. (2018). Application of metabolomics in Alzheimer’s disease. Frontiers in Neurology, 8(JAN), 1–20. https://doi.org/10.3389/fneur.2017.00719

Yin, P., Lehmann, R., & Xu, G. (2015). Effects of pre-analytical processes on blood samples used in metabolomics studies. In Analytical and Bioanalytical Chemistry (Vol. 407, Issue 17, pp. 4879–4892). https://doi.org/10.1007/s00216-015-8565-x

