## Supplementary Figures for "Deep metabolic profiling assessment of tissue extraction protocols for three model organisms"

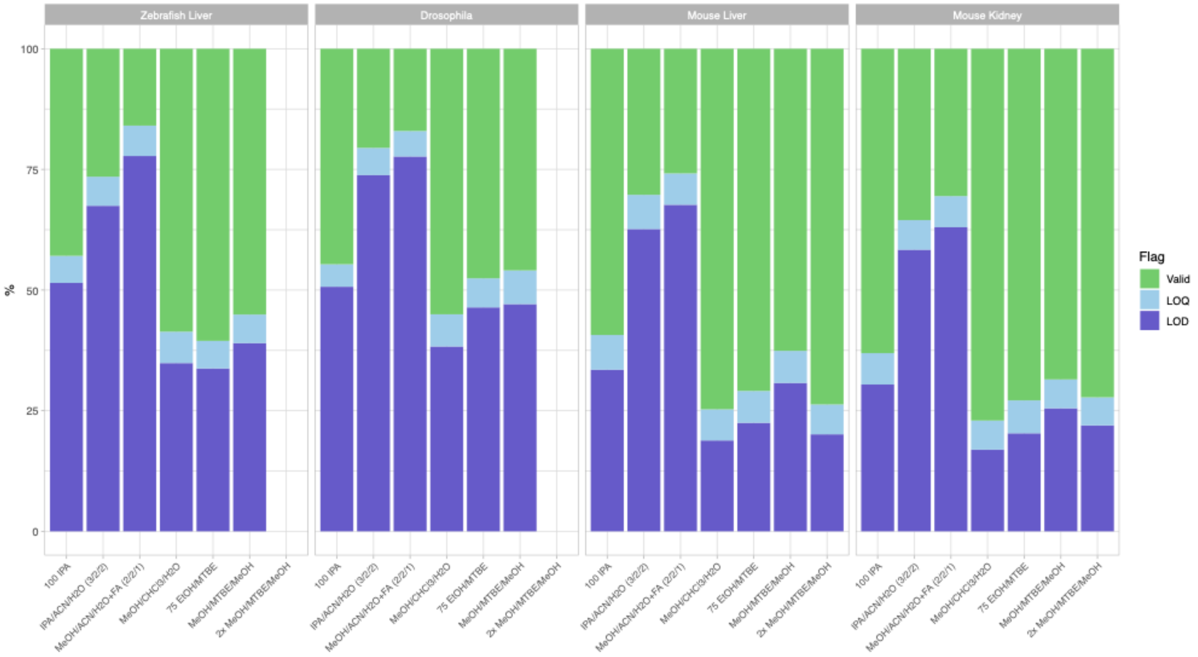


**Figure S1. Percentage of metabolite quantitation status across the seven extraction protocols per model organism and sample type.** The color shows all measurements that were performed at the limit of detection (LOD), lower/upper limit of quantification (LOQ) and within the quantitation range (valid).

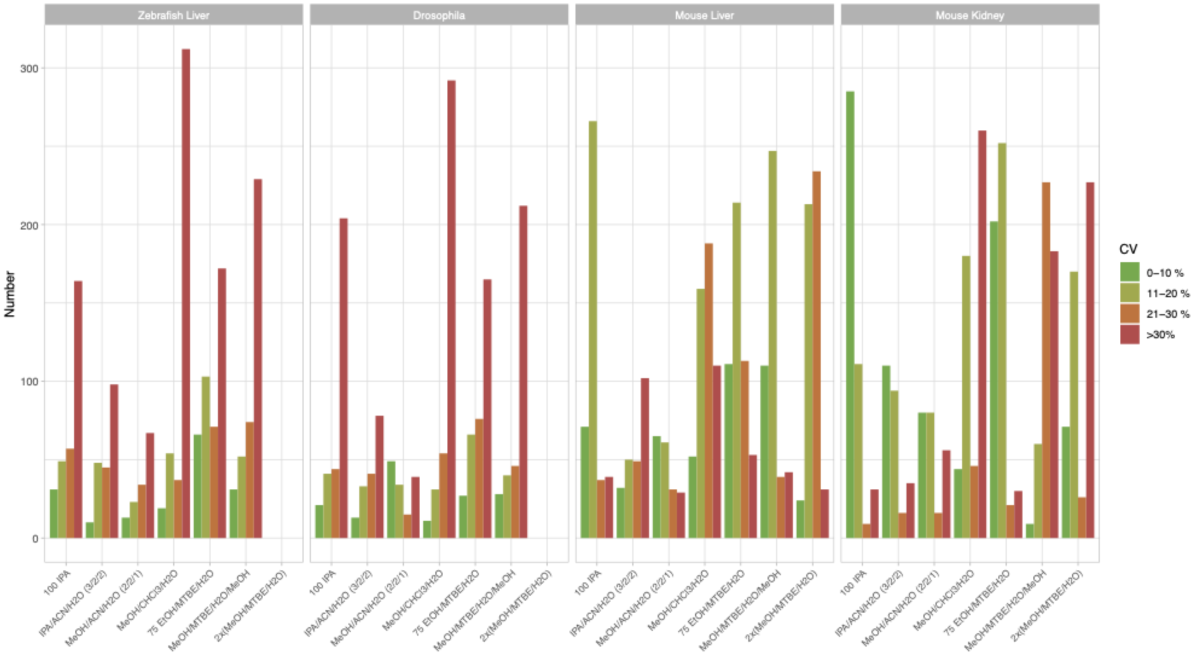

**Figure S2. Variability of metabolites across the seven extraction protocols per model organism and sample type.** Depicted is the CV% in different ranges and further rated using a color scheme from green (0-10%) to red (>30%). The number of metabolites accumulates to the total number of detectable metabolites in a given protocol.

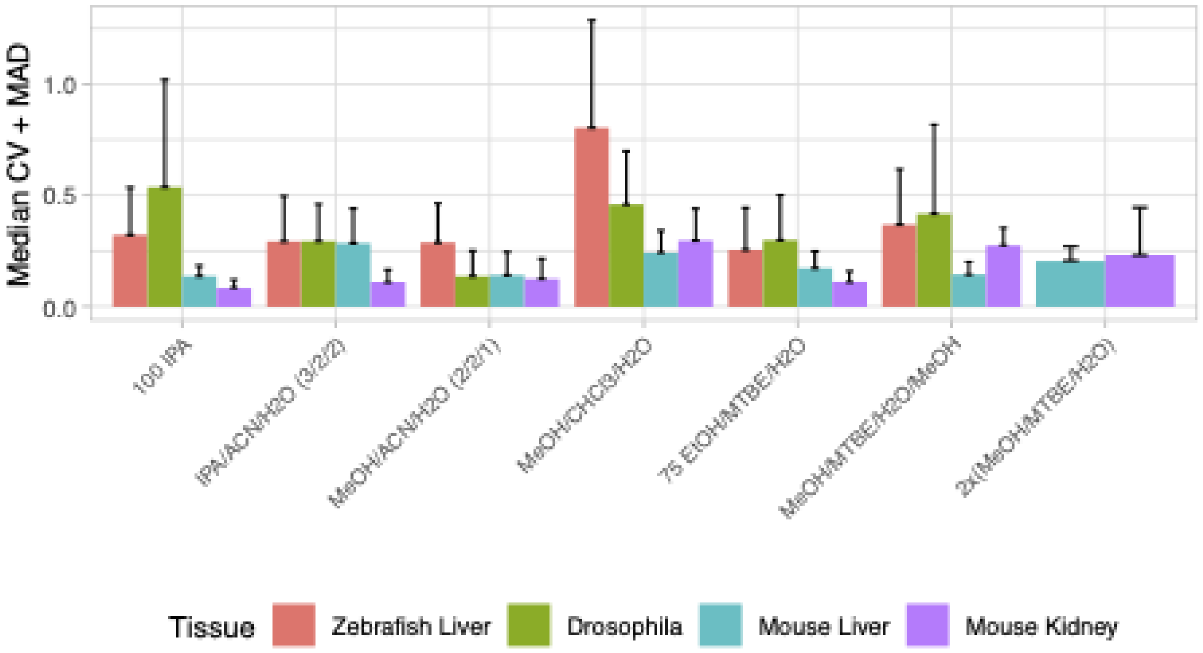


**Figure S3. Median and median absolute deviation (MAD) of the coefficient of variation (CV) across the seven extraction protocols.** Each bar represents a sample type across the different model organisms investigated. The order of extraction protocols follows Figure 1.

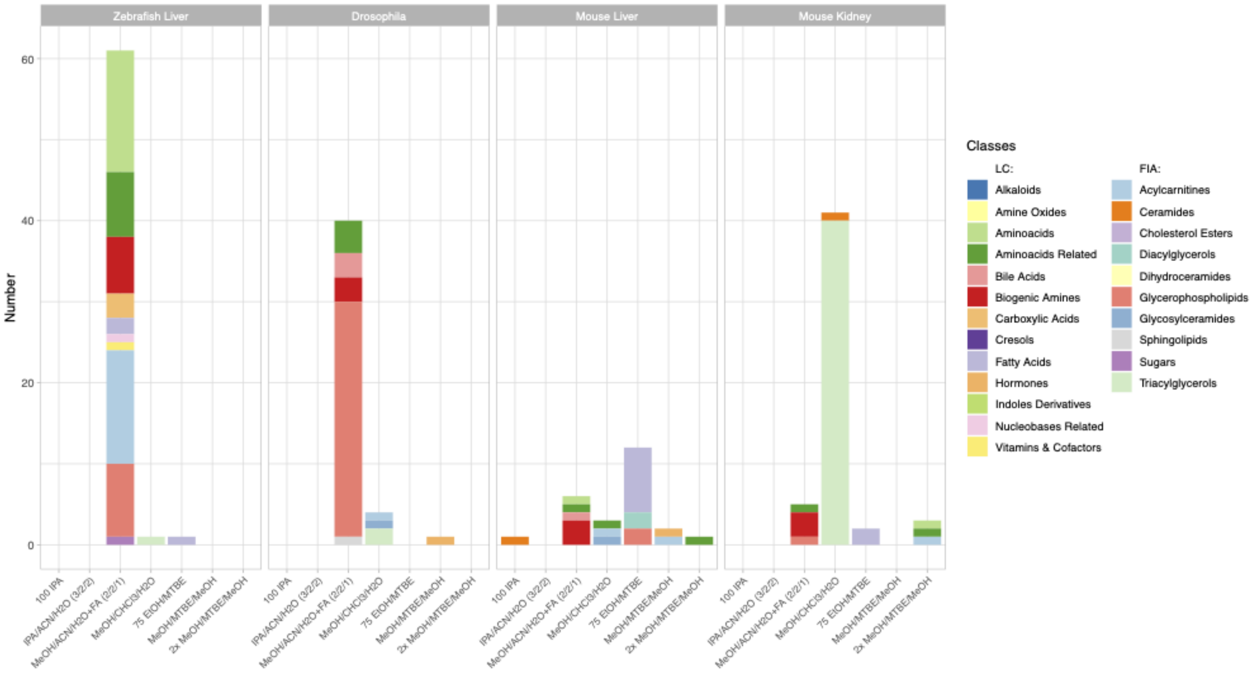


**Figure S4. Extraction protocols with exclusively high metabolite yield across the seven extraction protocols per model organism and sample type.** Indicated by color are the different metabolite classes. The legend is categorized between the LC-MS/MS and FIA-MS/MS measurements.
